# Light-controlled membrane remodeling in gel-fluid phase-separated giant vesicles using photoswitchable lipids

**DOI:** 10.64898/2026.04.20.719604

**Authors:** Tsu-Wang Sun, Elias Sabri, Mina Aleksanyan, Rumiana Dimova

## Abstract

Photoswitchable lipids enable optical control of membrane area, mechanics and phase behavior, offering a platform to study stimuli-responsive biomimetic systems. However, it remains unclear how the spatial organization of coexisting membrane phases governs photoinduced mechanical responses. Here, we incorporate the photoswitch azobenzene-phosphatidylcholine (azoPC) and dipalmitoyphosphatidylcholine (DPPC) into gel-fluid phase-separated giant unilamellar vesicles (GUVs) to probe how domain architecture controls light-driven deformation. We combine differential scanning calorimetry, temperature-controlled confocal microscopy, and a calibrated heating stage to quantify phase transitions and visualize domain dynamics. Dispersed domains produce global GUV crumpling upon UV-light-induced *trans*-to-*cis* isomerization of azoPC, whereas coarsened fluid domains locally confine deformation to budding regions of the GUVs; both responses are reversed by blue light. Temperature-controlled imaging reveals that the gel-fluid transition in GUVs is considerably broader than the calorimetric profile suggests, with coexisting phases detectable well above the calorimetry peak transition temperature. Well above the transition temperature, i.e. in the fully melted membrane, UV irradiation unexpectedly induces reversible nucleation of gel-like flower domains, consistent with an increased transition temperature in the *cis* azoPC state due to lipid packing incompatibility with DPPC. These results demonstrate that membrane domain architecture dictates the spatial distribution of photoinduced remodeling and that photoswitchable lipids can tune both membrane morphology and phase equilibria, enabling new strategies for stimuli-responsive synthetic cells and soft actuators.

## 1. Introduction

Biological membranes are complex and dynamic assemblies whose physical properties, such as fluidity, thickness, and elasticity, are tightly coupled to cellular function and disease. Their compositional diversity gives rise to lateral heterogeneity and thermodynamic phase behavior, including gel, liquid-ordered (Lo), and liquid-disordered (Ld) phases. Phase separation, driven by differences in lipid chain melting temperatures, represents a fundamental organizational principle of membranes and critically influences protein localization, signaling, and trafficking^1,2^. Through the interplay between lipid packing and membrane mechanics, phase behavior regulates processes ranging from adhesion and division to intracellular transport^3^.

In cellular plasma membranes, this heterogeneity is often conceptualized as lipid rafts, cholesterol- and sphingolipid-enriched domains that serve as platforms for organizing signaling components and facilitating protein-lipid interactions^4–6^. Although lipid heterogeneities in living cells are typically transient and nanoscopic, micron-scale phase separation has been observed in specific organisms such as yeast^7^. To systematically investigate the physical principles underlying membrane phase coexistence, model systems such as giant unilamellar vesicles (GUVs) provide a powerful and controllable platform^8,9^. In these systems, phase domains can be directly visualized by fluorescence microscopy using probes that preferentially partition into specific lipid environments, for example, enriching in fluid domains while being excluded from gel phases^10^. Similar to lipids, membrane-associated proteins and peptides exhibit preferential partitioning depending on their structural and chemical compatibility with local membrane order, thereby coupling domain organization to functional regulation^4,11,12^.

Gel-fluid phase coexistence profoundly alters the mechanical properties of membranes. The presence of distinct gel and fluid phases drastically affects membrane elasticity^13,14^ with ordered and gel phases generally exhibiting higher bending moduli than disordered ones^15,16^. This heterogeneity affects not only elasticity but also permeability, as interfaces between coexisting phases can act as preferential pathways for ion or solute transport^17^. Moreover, the coupling between in-plane molecular order and topological constraints can drive dramatic morphological transformations, including shape remodeling and vesicle fission under non-equilibrium conditions such as thermal gradients^18,19^. These phenomena highlight the central role of phase behavior in governing membrane architecture and responsiveness.

Harnessing phase coexistence in model membranes enables the rational design of systems with tunable mechanical and transport properties. Phase-separated vesicles have been explored for controlled drug release, where permeability can be selectively modulated through domain organization, and for biosensing applications in which changes in domain morphology serve as responsive readouts^20^. Their integration as functional elements in soft robotics, where membrane deformations generate directional motion or force^21–23^ further illustrates the potential of membrane phase behavior as an engineering handle. Realizing the full potential of these systems, however, requires external stimuli capable of dynamically and reversibly modulating membrane properties with high spatiotemporal precision.

Molecular photoswitches provide a powerful strategy for achieving noninvasive, reversible control over soft matter systems using light^24^. Among these, azobenzene-phosphatidylcholine (azoPC), a synthetic lipid with an azobenzene moiety within one of the lipid tails (Fig. 1A), has emerged as a versatile component for synthetic membrane systems and even for applications in living cells. Upon UV-A (365 nm) irradiation, azoPC undergoes *trans*-to-*cis* isomerization, which can be reversed with blue light (465 nm), enabling dynamic modulation of membrane properties. In homogeneous fluid membranes, azoPC photoisomerization has been shown to induce substantial membrane area increases (up to ∼25%) and dramatic softening, reflected in up to an order-of-magnitude reduction in bending rigidity^25^. It further leads to decreased membrane thickness and enhanced fluidity^26,27^, and can promote reversible morphological transformations such as deformation^28^, budding ^25^, and endocytosis-like responses ^29^. Beyond shape changes, light-triggered azobenzene isomerization has been linked to altered membrane permeability and transient pore formation, enabling controlled solute transport^30^. In liquid-liquid (Lo/Ld) phase-separated ternary lipid mixtures, azoPC has been shown to reversibly reorganize membrane domains, with UV irradiation converting liquid-ordered regions into liquid-disordered ones^5,31^. Together, these studies establish azobenzene lipids as precise tools for spatiotemporal control of membrane structure and function, with demonstrated impact on protein activity^32–34^, and cellular morphology in living systems^35,36^.

**Figure 1.**
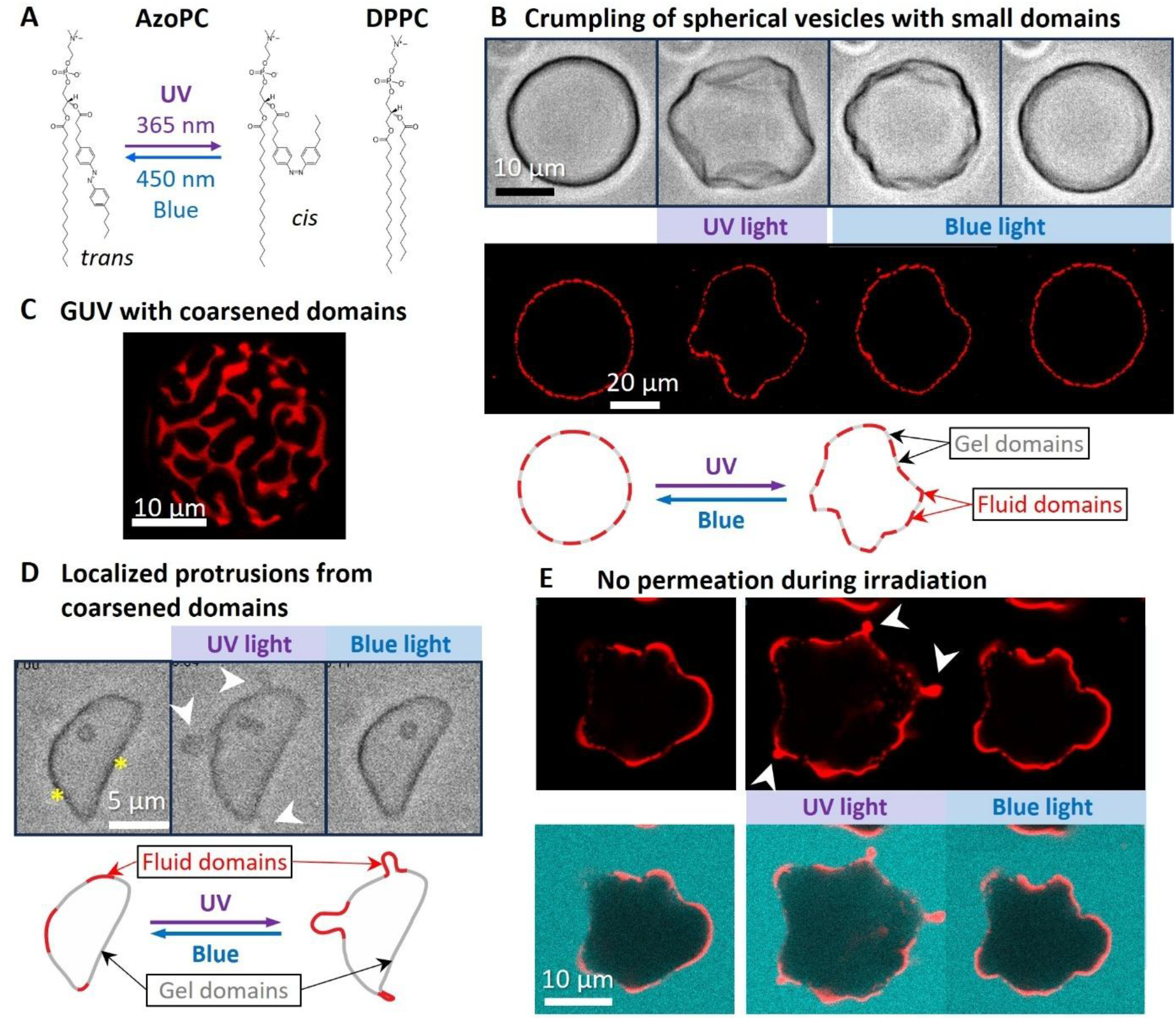
Light-induced crumpling and localized budding in phase-separated azoPC:DPPC 1:1 vesicles. (A) Chemical structures of the used lipids. Left: azoPC in its *trans* and *cis* configurations, interconverted by UV (365 nm) and blue (here, 460 nm) light; right: DPPC. (B) Light-induced crumpling of GUVs with small dispersed domains. Top: Brightfield images of a GUV showing global membrane crumpling upon UV irradiation, fully reversed under blue light, see Movie S1. Middle: Confocal cross sections of another GUV under UV irradiation illustrating overall deformation: the rigid gel domains resist expansion while fluid regions allow local distortion. Bottom: cartoon summarizing the overall crumpling of GUVs with disperse small domains. (C) Confocal z-stack projection of a half of a GUV after slow stepwise cooling leading to domain coarsening. Dark, branched domains correspond to DPPC-rich gel regions excluding the fluorescent dye (0.1 mol% Atto-647N-DOPE), while bright regions are azoPC-rich fluid domains. (D) Spatially confined budding in vesicles with coarsened gel domains. Top: Brightfield image sequence of a deflated azoPC:DPPC GUV with coarsened domains: flat facets (asterisks) correspond to rigid DPPC-rich gel regions that remain practically immobile under UV irradiation (middle), while curved fluid segments between them protrude as localized buds (arrowheads). Buds retract fully upon blue light irradiation (right). A similar example GUV response is shown in Movie S2. Bottom: cartoons illustrating the localized budding response when domains are coarsened contrasting the overall crumpling of vesicles with disperse domains as shown in panel B. (E) No permeation during isomerization. Confocal image sequence of a vesicle with coarsened domains labeled with 0.1 mol% Atto-647N-DOPE (red) and sulforhodamine B (SRB, cyan) added to the external solution (top: membrane channel only, bottom: merged signal). Localized budding of the coarsened fluid domains (arrowheads) is observed under UV irradiation and fully reversed under blue light, see also Movie S3. The absence of SRB signal inside the vesicle throughout confirms membrane integrity.

Despite this progress, the interplay between photoswitchable lipids and gel-fluid phase coexistence remains largely unexplored. Gel-fluid systems differ fundamentally from the Lo/Ld mixtures studied previously: the mechanical contrast between phases is far more pronounced, with gel domains exhibiting bending moduli orders of magnitude higher than coexisting fluid regions^16^ and line tensions at gel-fluid boundaries are substantially higher^13^. It is therefore unclear how light-induced conformational changes propagate within such mechanically heterogeneous membranes, and whether the spatial segregation of elastic properties confines or redirects photoinduced deformation. Here, we introduce azoPC into gel-fluid phase-separated GUVs to investigate how *trans-cis* isomerization influences domain stability, spatial confinement of deformation, and membrane remodeling dynamics. By combining controlled temperature variation with UV and blue light irradiation, we systematically examine how domain size, organization, and melting behavior modulate the membrane response to light. This approach allows us to dissect how pre-existing mechanical heterogeneity governs the extent and localization of photoinduced transformations, revealing new principles for dynamically controlling complex membrane architectures.

## 2. Materials and Methods

### 2.1 Materials

The lipids 1-stearoyl-2-[(E)-4-(4-((4-butylphenyl)diazenyl)phenyl)butanoyl]-*sn*-glycero-3-phosphocholine (azoPC), 1-palmitoyl-2-oleoyl-*sn*-glycero-3-phosphocholine (POPC) and 1,2-dipalmitoyl-*sn*-glycero-3-phosphocholine (DPPC) were purchased from Avanti Polar Lipids (Alabaster, AL). Glucose, sucrose, and the fluorescent dye 1,2-dioleoyl-*sn*-glycero-3-phosphoethanolamine labeled with Atto 647N (Atto-647N-DOPE) were purchased from Sigma-Aldrich (St. Louis, MO, USA). Sulforhodamine B was purchased from Thermo Fischer (Germany). Chloroform (LiChrosolv®) was purchased from Merck KGaA, Germany. Lipids (and 0.1 mol% of Atto-647N-DOPE) were dissolved in chloroform at typical concentration of 4 mM, and the stock solutions were stored at -20 °C until use. Sulforhodamine B for membrane leakage was prepared as a stock concentration 220 μM.

### 2.2 Vesicle preparation

The electroformation of GUVs was conducted in an oven at 65°C to ensure the temperature was above the melting point of the lipids. An aliquot of 12-30 μL of the lipid solution in chloroform was spread onto two 50 mm × 50 mm indium-tin-oxide (ITO) coated glass slides (Präzisions Glas & Optik GmbH, Germany). The slides were then clamped together using a 2 mm Teflon spacer and forming a chamber which was then filled with 100 mM sucrose solution (resulting in roughly 25 μM lipid in the final GUV solution; note that for calorimetry measurements, we doubled this concentration to roughly 55 μM to increase the signal-to-noise ratio). The chamber was placed in the oven for 5 minutes before the application of a sinusoidal alternating current (AC) field at 10 Hz and 1 V root-mean-square (RMS) for 1.5 hours. Following electroswelling, the oven was turned off, and the system was allowed to cool gradually to room temperature over 2 hours. A 0.75 mm Type K Miniature Thermocouple (TC Direct) inserted into the sample recorded a cooling rate of 0.3 K/min (see Fig. S1 in the Supporting information, SI).

Multilamellar vesicles (MLVs) were prepared using the freeze-thaw method following Ref. ^37^. Lipid solutions in chloroform were first dried in a glass vial under a stream of nitrogen, followed by desiccation for 2 hours. A sucrose solution (100 mM) was warmed up to 65°C and added to achieve a final lipid concentration of 1 mg/mL, and the mixture was vortexed for 30 seconds. The sample was then subjected to three cycles of heating in a 65°C water bath, cooling on ice, and vortexing.

### 2.3 Differential scanning calorimetry (DSC)

DSC measurements were performed using a MicroCal VP-DSC system (MicroCal, USA). Both MLVs and the reference sucrose solution (100 mM) were degassed (ThermoVac, MicroCal, USA) for 5 minutes before being introduced into the measurement and reference chambers with volumes of approximately 0.5 mL. The scanning mode was set to a rate of 1 K/min. Data analysis was conducted using the baseline subtraction functions in Origin 2023 (OriginLab, USA).

### 2.4 Microscopy and vesicle contour analysis

GUVs were imaged using an inverted Leica TCS SP8 scanning confocal microscope (Leica, Germany) equipped with a 63×, 1.2 Numerical Aperture (NA) water immersion HC PL APO CS2 objective, and an HC PL FLUOTAR L 40×, 0.60 NA objective. Atto-647N-DOPE was excited with a Helium-Neon (HeNe) laser at 633 nm (0.5% laser intensity) and the emission signal was collected in the range 643–700 nm with photomultiplier detector. Scanning was performed at 600 Hz in bidirectional mode, with the pinhole size adjusted to 1-4 Airy units (AU), where 1 AU was used for stacked images used for 3D reconstruction, and larger AUs were employed for observing irregular deformations.

Membrane dynamics were recorded using an inverted Zeiss Axio Observer D1 microscope (Carl Zeiss, Germany) equipped with a 63×, 1.2 NA water immersion C-Apochromat objective and a Pco.Edge sCMOS camera (PCO Imaging, Germany), operating at an acquisition rate of 20 frames per second. The observation chamber consisted of two coverslips and a Press-to-Seal silicone isolator (Invitrogen, Eugene OR, USA) as a spacer. The vesicles were recorded under bright field.

To quantify the degree of GUV deformation, we utilized a contour-tracking algorithm developed in Ref.^38^. Briefly, the algorithm assigns a set of interpolation points to the GUV perimeter, where the number of points (*N*) in the contour scales with vesicle size. By monitoring the coordinates of these points over time, we quantify the degree of shape deviation from a spherical geometry based on a weighed cumulated relative displacement *δ* of contour points from their initial coordinates, taken as the onset of UV irradiation, via the expression

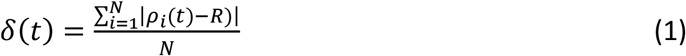

where *ρ*_*i*_(*t*) is the distance of the *i*-th point to the center of the GUV (defined via the averaged horizontal and vertical coordinates of all points on the contour) at time *t* and *R* is the initial average distance of contour points from the center of the vesicle.

### 2.5 Light-induced isomerization

Two distinct light-emitting diodes (LED) were utilized to precisely control the isomerization of azoPC lipids: a high-power UV-LED system and a versatile fiber-coupled LED system.

For broad-field UV irradiation, a home-built UV-LED system was employed. This unit comprised a cooling metal plate and an LED (Star-UV365-01-00-00, Roschwege, Germany) with a maximum power output of 170 mW. The UV-LED unit was positioned directly onto the sample observation chamber during experiments to ensure uniform illumination (see Fig. S2a in the SI).

For wavelength-specific illumination, a fiber-coupled LED system (Doric, Canada) was used. This system featured two distinct light sources: a UV LED (365 nm) and a blue LED (450 nm). Light intensity was controlled via an integrated driver, which offered both manual adjustment via a control panel and software-based regulation through a connected laptop. The optical fiber (960 μm, 0.63 NA) from this system was precisely affixed alongside the microscope objective using a cantilever arm to direct irradiation onto the sample (see SI Fig. S2b). The maximum power outputs, measured at the sample plane, were determined using a LaserMate 10 sensor and console (Coherent, USA). The UV light (365 nm) delivered a maximum power of 75 mW, while the blue light (450 nm) delivered 183 mW.

### 2.6 Temperature control

Temperature control of the GUV samples under the microscope was precisely maintained using a custom-built electrical resistance heater with an observation chamber attached underneath and mounted on the confocal microscope. The heater consisted of two ITO-coated glass slides (25 mm × 50 mm) with adhesive copper tape fixed along their sides. These slides were clamped together with their conductive surfaces facing each other, forming the heating element, see SI Figs. S3 (also introduced in more detail later in the text and Fig. 4A). To maintain precise temperature control, the copper tapes were connected to a direct current (DC) generator (Programmable laboratory power supply unit HCS 3202, Manson, Hong Kong), which was operated via a laptop and regulated by a custom-developed Proportional–Integral–Derivative (PID) controller (Multi Control program, in-house developed).

Temperature feedback was established using a 0.5 mm diameter Mineral Insulated Thermocouple (TC Direct, UK), which was directly inserted into the sample chamber. This thermocouple was connected to a USB-TC01 temperature measurement device (National Instruments, USA) to provide real-time temperature readings to the PID controller. To prevent overheating, the maximum current supplied to the heater was limited to 1 A.

To maintain thermal stability and prevent heat dissipation through the objective, an objective heater (Tempcontrol mini, PeCon GmbH, Germany) was attached to the objective and synchronized with the PID controller for measurements between 30°C and 40°C, while remaining fixed at 40°C for all sample temperatures exceeding that range.

Prior to experiments, the thermocouples were calibrated by immersion in boiling and ice water. These measurements were cross-referenced against a Pt1000 probe connected to a Testo 735 temperature measuring device (Testo, Germany) to ensure accuracy. For experiments involving temperature simulations (assessing thermal distribution), calibration measurements were also performed by attaching thermocouples directly to the coverslip of the observation chamber and on top of the ITO heater using thermal paste (Peltier contact gel-silver filled, innovatek, Germany) and heat-resistant tape to minimize heat dissipation and ensure accurate local temperature readings.

### 2.7 Numerical simulations

Numerical simulations were performed using the finite element method software COMSOL Multiphysics (COMSOL Multiphysics® v. 5.2. www.comsol.com. COMSOL AB, Stockholm, Sweden). The model geometry (Fig. S4) was discretized using triangular elements with refined boundary layers at fluid interfaces. Heat transfer and fluid flow were solved in a coupled manner: in the fluid domains (water and air), the heat transfer equation was coupled to the incompressible Navier-Stokes equations within the Boussinesq approximation to account for buoyancy-driven convection, while in solid domains (glass, rubber, steel, and ITO) heat conduction was solved. Dirichlet temperature boundary conditions were imposed at the ITO interface and on the objective, which were adjusted to match experimentally measured temperatures at defined probe positions. Full details of the governing equations, boundary conditions, mesh statistics, and material parameters are provided in the SI (see also Fig. S4 and Table S1).

## 3. Results and Discussion

We begin by demonstrating that incorporating azoPC into gel-fluid phase-separated membranes gives rise to a qualitatively distinct mechanical response to light, manifested as pronounced crumpling rather than the smooth budding observed in homogeneous fluid vesicles. We then characterize how this response depends on UV intensity and establish its kinetic signatures. Finally, we determine the thermotropic phase behavior of the binary azoPC:DPPC system by calorimetry and temperature-controlled microscopy, providing the thermodynamic framework needed to interpret the light-driven phenomena described in subsequent sections.

### 3.1. Light-induced crumpling in gel-fluid phase-separated vesicles

To investigate how membrane heterogeneity influences photoswitchable deformation, we incorporated DPPC, a gel-phase lipid with a melting temperature of 41°C^39^, into azoPC-containing membranes, see Fig. 1A for lipid structures. GUVs were prepared as cell-sized model membranes^8,9^, enabling direct visualization of phase separation^40^ and light-induced remodeling^25^.

We focused on equimolar azoPC:DPPC mixtures to promote extended domain formation and maximize mechanical contrast between gel and fluid regions. The melting temperature (T_m_) of azoPC has not been reported to our knowledge. Although its stearoyl-containing tail might superficially suggest behavior similar to DSPC (1,2-distearoyl-*sn*-glycero-3-phosphocholine, T_m_ = 55°C^41^), the presence of a saturated chain alone does not dictate the phase behavior: SOPC (1-stearoyl-2-oleoyl-*sn*-glycero-3-phosphocholine), which also carries a stearoyl chain but also one unsaturated tail, has a T_m_ of 6°C^42^. More directly, pure azoPC GUVs display thermal shape fluctuations and bending rigidities characteristic of fluid membranes at room temperature^25^, indicating that the T_m_ of azoPC lies below 23°C.

Binary mixtures of gel- and fluid-phase lipids, such as DPPC:POPC or DPPC:SOPC, are well known to exhibit gel-fluid phase coexistence at room temperature, typically characterized by micron-sized gel domains embedded in a fluid matrix depending on composition and thermal history. By analogy, we expect equimolar DPPC-azoPC membranes at room temperature to phase-separate into DPPC-rich gel domains and azoPC-rich fluid domains.

After electroformation at 60°C, the GUVs were cooled in the oven gradually (∼0.3 K/min; Fig. S1 in the supporting information), occasionally with additional waiting steps at fixed temperatures, to promote domain coarsening. Slow cooling is known to enhance domain growth by allowing lipid demixing to approach equilibrium and reducing kinetic trapping of small domains^43^ as we will discuss below. The resulting vesicles were examined by confocal microscopy.

Fluorescent labeling with 0.1 mol% Atto-647N-DOPE revealed branched, dye-excluding domains (Fig. 1B,C), consistent with gel-phase DPPC regions excluding the fluorescent lipid. We assign the fluorescent regions to azoPC-rich fluid domains and the dark, finger-like regions to DPPC-rich gel domains. Faster cooling resulted in smaller, more dispersed domains (Fig. 1B, lower panel), compared to larger, coarsened domains in vesicles where cooling was done slowly in a stepwise manner (Fig. 1C), confirming the sensitivity of domain size to thermal history.

GUVs were irradiated with UV or blue light using a fiber-coupled LED device with precise intensity control, typically set at 500 mA, corresponding to approximately 75 mW of power at the sample (Fig. S5). As shown previously^25^, fluid azoPC-doped vesicles exhibit membrane area expansion stored in transient buds under UV irradiation, yielding a floppy morphology that reverses under blue light (Fig. S6). Pure DPPC vesicles show no morphological response to UV light.

In marked contrast to fluid azoPC-doped vesicles, equimolar azoPC:DPPC vesicles with dispersed domains respond to UV light with pronounced crumpling and twisting deformations (Fig. 1B). Rather than forming large buds leading to homogeneous enlargement, as observed in homogeneous fluid membranes (Fig. S6), the area expansion appears spatially constrained. The softer azoPC-rich regions undergo local distortion, while the rigid DPPC-rich domains act as mechanical scaffolds that resist uniform expansion. The resulting morphology reflects mechanical frustration between expanding fluid domains and mechanically stiff gel regions (Fig. 1B). Upon blue light irradiation, the vesicles fully recover their initial morphology (Fig. 1B; see also Movie S1), demonstrating reversible, light-controlled mechanical remodeling in a heterogeneous membrane.

The mechanical response to UV irradiation depends critically on domain size and connectivity, which in turn are determined by thermal history. In the dispersed-domain regime described above, UV irradiation produces global crumpling distributed across the entire vesicle surface (Fig. 1B). However, when domains are coarsened by slow passage through the melting transition (as discussed below), a qualitatively different response emerges: deformation becomes spatially confined to the fluid regions, while the gel domains act as a rigid, unresponsive scaffold (Fig. 1D).

Coarsened gel domains in deflated vesicles produce characteristic flat facets on the vesicle surface. These are planar segments whose presence reflects the high stiffness of the membrane in the gel domains^15^, which resist the curvature that the fluid membrane would otherwise adopt^14,44^. This faceted morphology is visible in brightfield imaging (Fig. 1D, asterisks).

Upon UV irradiation, the faceted regions remain entirely immobile, while the curved fluid segments between them, which are enriched in azoPC due to the lower bending rigidity of the fluid phase favoring higher-curvature regions, protrude as localized buds (arrowheads in Fig. 1D,E, and sketch in D; see also Movies S2 and S3). These buds can protrude outward and inward imposed by the constant volume constraint. The spatial confinement of the overall vesicle deformation arises because the gel domains are mechanically inextensible. The excess area generated by *trans*-to-*cis* photoisomerization is localized predominantly into the fluid azoPC-rich islands, which store it as outward or inward protrusions.

The localized budding is fully reversed upon blue light irradiation, with the vesicle returning to its pre-irradiation faceted morphology (Fig. 1D,E; Movies S2, S3), confirming that the process is driven by photoisomerization and not by irreversible membrane remodeling.

To verify membrane integrity during this repeated deformation, sulforhodamine B (SRB) was added to the outer solution (to a final concentration of ∼50 µM) prior to irradiation. No SRB signal was detected inside the vesicles after more than 10 cycles of UV (5 s each) followed by blue light (5 s) irradiation, confirming that no membrane leakage occurs during budding (Fig. 1E, merged channel, Movie S3).

Together, the two regimes of (i) global crumpling with dispersed domains (Fig. 1B) and (ii) localized budding with coarsened domains (Fig. 1D,E) demonstrate that domain size functions as a structural programmable parameter that determines how photoswitchable area expansion is distributed across the membrane. Dispersed domains below the percolation threshold allow global mechanical frustration; coarsened continuous (percolating) domains above it focus deformation into spatially defined fluid islands. This principle, namely, using gel domain architecture to direct the localization of light-driven membrane remodeling, offers a strategy for achieving spatially programmable morphological control in synthetic membrane systems.

### 3.2. Deformation dynamics depend on UV intensity

To quantify deformation kinetics, we systematically varied UV intensity while remaining well below previously reported damage thresholds (280–440 mW/cm^2^) associated with membrane defects^45^. Here, intensities ranged from 10 to 75 mW/cm^2^.

The crumpling response in GUVs with small dispersed domains was highly reproducible across multiple irradiation cycles of the same vesicle, with morphology returning to its original state after each blue-light exposure (Fig. S7; 13 cycles). The absence of residual deformation suggests that no irreversible photodamage or photooxidation occurs under our conditions. The fully reversible shape recovery also indicates that membrane permeability remains unchanged within experimental resolution; no leakage events were observed during repeated cycling.

To quantify shape changes, we tracked vesicle contours using a previously developed algorithm^38^, which detects the vesicle outline, and computed the average contour displacement relative to the initial shape (see Eq. 1). This metric captures deformation independently of direction, overcoming limitations of simple radius or perimeter measurements.

Representative time traces for a single vesicle at different UV intensities are shown in Fig. 2A. The displacement amplitude increases and the response accelerates with increasing UV intensity. The temporal evolution was fitted with an exponential model, *δ*(*t*) *= δ*_0_ *+ A*_1_(1 − *e*^−*t/τ*^) from which a characteristic response time *τ* was extracted for each trial (fits shown in Fig. 2A).

**Figure 2.**
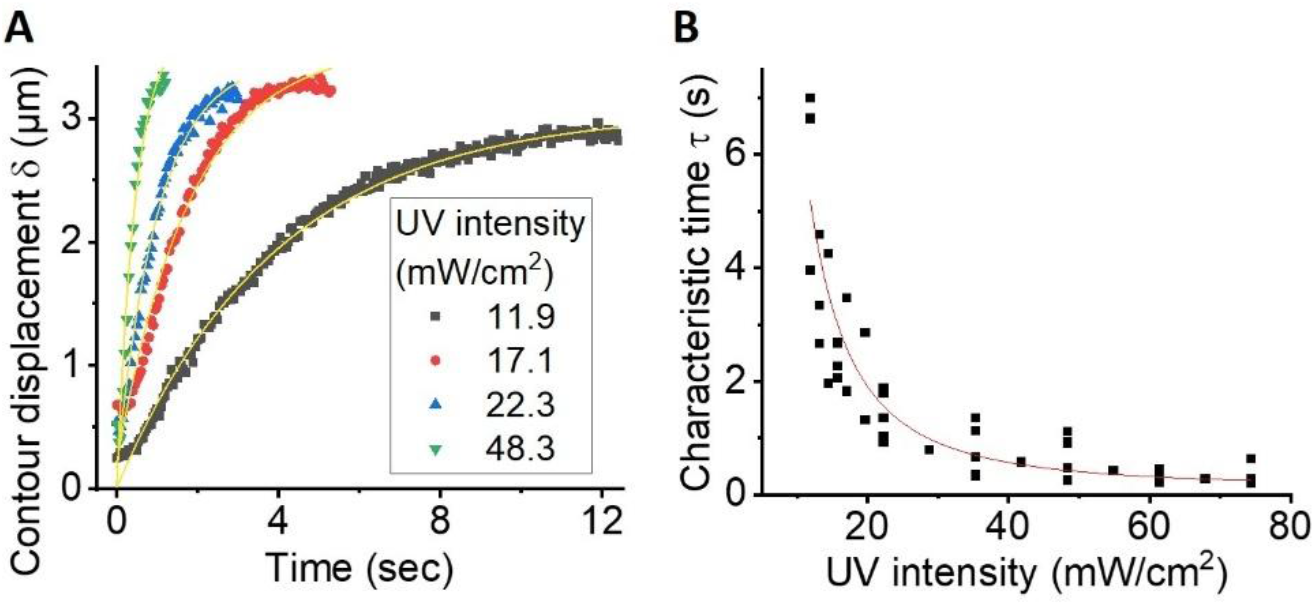
UV intensity controls deformation kinetics in azoPC-DPPC vesicles (1:1 molar ratio). (A) Time dependence of the relative contour displacement *δ* (defined in Eq. 1) under different UV intensities (indicated in the legend) applied to the same GUV. Solid lines (yellow) are single-exponential fits: *δ = δ*_0_ *+ A*_1_(1 − *e*^−*t/τ*^).The vesicle diameter was ∼30 µm. (B) The characteristic response time *τ* as a function of UV light intensity plateaus at higher UV intensity, consistent with saturation of the photoisomerization process. Data from 4 vesicles. The red curve is an exponential decay fit 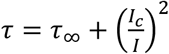 with *τ* = 0.12 s and *I*_*c*_= 26.7 mW/cm^2^ illustrating the saturation behavior.

Across vesicles of varying sizes (radius between 15 and 30 µm), *τ* decreased with increasing UV intensity (Fig. 2B). Above approximately 27 mW/cm^2^, the response times approached a plateau, suggesting saturation of the photoisomerization process. This behavior is consistent with a steady-state balance between *trans-*to-*cis* conversion under UV illumination and spontaneous thermal back-isomerization.

The scatter in *τ* at fixed intensity likely reflects vesicle-to-vesicle differences in membrane tension, since GUV electroformation does not allow precise tension control, as well as variations in domain size and connectivity.

These results establish UV intensity as a tunable parameter controlling deformation kinetics. To understand how the thermotropic phase behavior of the binary system underlies this mechanical response, we next characterize its transition temperature.

### 3.3. Thermotropic behavior of azoPC-DPPC mixtures

To determine the phase transition temperature of azoPC-containing membranes, we performed differential scanning calorimetry (DSC) on MLV suspensions (∼1 mg/mL lipid) and compared them to more dilute GUV suspensions at approximately 20-fold lower lipid concentration. To increase the signal-to-noise ratio of the GUV measurements, the amount of lipid deposited on the ITO-coated glasses during electroformation was doubled relative to the preparations used for microscopy observations; note that even at this higher deposition, the lower total lipid content of GUV suspensions relative to MLV suspensions results in a low signal-to-noise ratio.

Pure DPPC samples exhibited a sharp main transition at ∼41°C (Fig. S8), consistent with literature values^39^. Pure azoPC membranes exhibited no detectable transition between 20 and 60°C, consistent with a fluid state throughout this temperature range.

Equimolar azoPC:DPPC mixtures displayed a prominent peak centered at 31.9°C (Fig. 3), superimposed on a broad baseline with a half-width of approximately 3°C. This curve shape indicates partial cooperativity: the sharp peak likely reflects melting of DPPC-rich domains, while the broad base is characteristic of extended gel-fluid coexistence in non-ideal binary mixtures^46^. The transition temperature of 31.9°C, which is approximately 10°C below the T_m_ of pure DPPC, reflects the fluidizing influence of azoPC on the mixed membrane.

**Figure 3.**
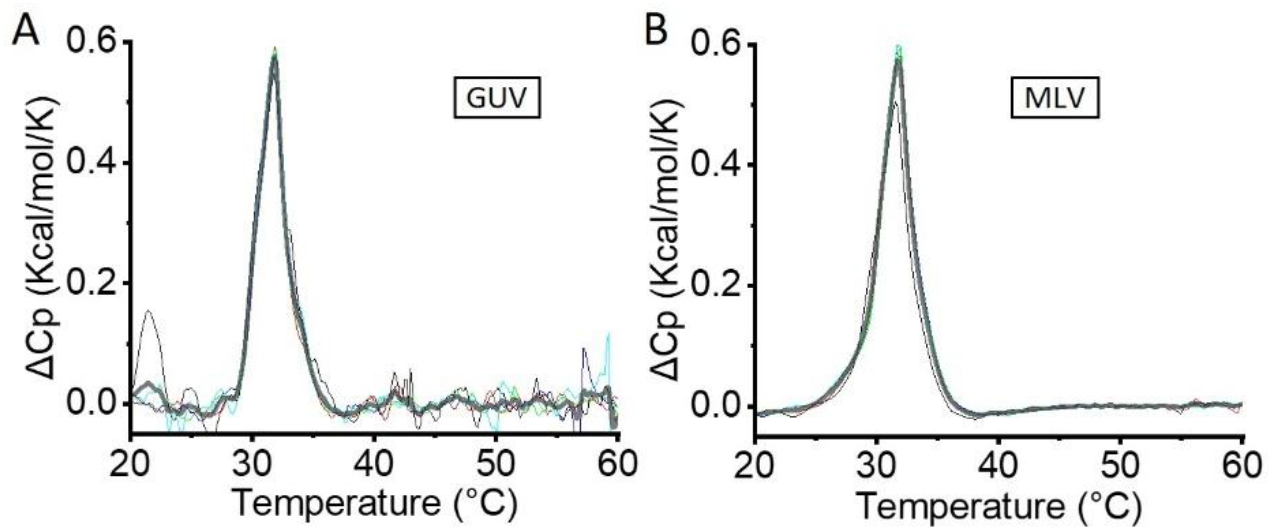
Differential scanning calorimetry of 1:1 azoPC:DPPC mixtures. DSC traces of (A) GUV suspension (55 µM total lipid; 43 µg/mL) prepared with doubled lipid deposition relative to microscopy samples to improve signal-to-noise and (B) MLV suspension (1 mg/mL); both samples were prepared in 100 mM sucrose. Heating rate: 1 K/min. The main transition temperature is ∼31.9 °C, approximately 9 °C below the T_m_ of pure DPPC (Fig. S8). Note that the GUV profile may contain contributions from non-unilamellar material and is expected to represent a lower bound on the true breadth of the gel-fluid transition in isolated GUV bilayers (see main text). Colored curves represent five consecutive heating scans, with the averaged profile shown as a thick grey line.

We note, however, that the interpretation of the GUV DSC signal warrants caution. The increased lipid deposition during electroformation used to improve signal-to-noise may introduce a contribution from non-GUV material, including multilamellar vesicles or lipid aggregates that did not fully form into unilamellar structures, and we cannot exclude the extent to which these populations influence the measured profile. For genuinely unilamellar GUV bilayers, the transition would be expected to be broader than the MLV profile, for two reasons: First, the higher cooperativity of MLV transitions arises from interlamellar coupling between stacked bilayers^37^, which is absent in isolated GUVs. Second, the reduced cooperative unit size and higher effective impurity levels in GUV samples^47^ could further broaden the thermotropic response. Indeed, LAURDAN-based two-photon fluorescence microscopy measurements of the gel-to-liquid crystalline transition in single-component GUVs revealed a broad transition range and coexisting membrane regions of distinct generalized polarization over an extended temperature window^48^. Considering these effects, the true gel-fluid transition in the azoPC:DPPC GUVs is likely broader than what the DSC curves suggest, with gel-fluid coexistence potentially detectable more than 10 K above and below the peak transition temperature of 31.9 °C (as shown previously in LAURDAN-based measurements on GUVs^48^). This is also consistent with the microscopy observations described in section 3.5, where gel domains persist at sample temperatures considerably above the calorimetric transition.

Interestingly, despite the expectation that interconnected gel domains might form a mechanically continuous scaffold constraining deformation, microscopy revealed relatively dispersed dendritic domains even after slow oven cooling (Fig. 1C). Attempts to further increase domain size by reheating above the transition temperature and re-cooling at the same rate (0.3 K/min) did not significantly alter domain morphology. This is consistent with domain connectivity remaining below the percolation threshold required for spatially confined deformation, and provides a thermodynamic rationale for the global crumpling morphology observed in Fig. 1B, rather than the localized buckling that would be expected for a fully percolating gel network. The latter regime, i.e. spatially confined, localized budding, could be accessed only upon slow passage through the melting transition with prolonged equilibration at temperatures just above the first microscopic appearance of domains, allowing sufficient time for domain coarsening and coalescence to build a mechanically continuous gel scaffold.

These thermodynamic measurements provide the framework for the temperature-controlled microscopy experiments described in the following section, in which we directly visualize domain melting in GUVs and its coupling to light-driven membrane remodeling.

### 3.4. Temperature gradients and numerical calibration of a custom heating chamber for real-time confocal microscopy observations

To observe temperature-dependent domain reorganization, we require a device that maintains optical accessibility for confocal microscopy while heating the sample. We developed a novel custom-made heating device, consisting of ITO glasses and a DC generator, regulated via a PID control program with feedback from the temperature measurements inside the sample. This setup allowed us to maintain optical transparency while heating the samples, enabling real-time imaging under confocal microscopy (Fig. 4A, section 2.6 in the Materials and Methods).

**Figure 4.**
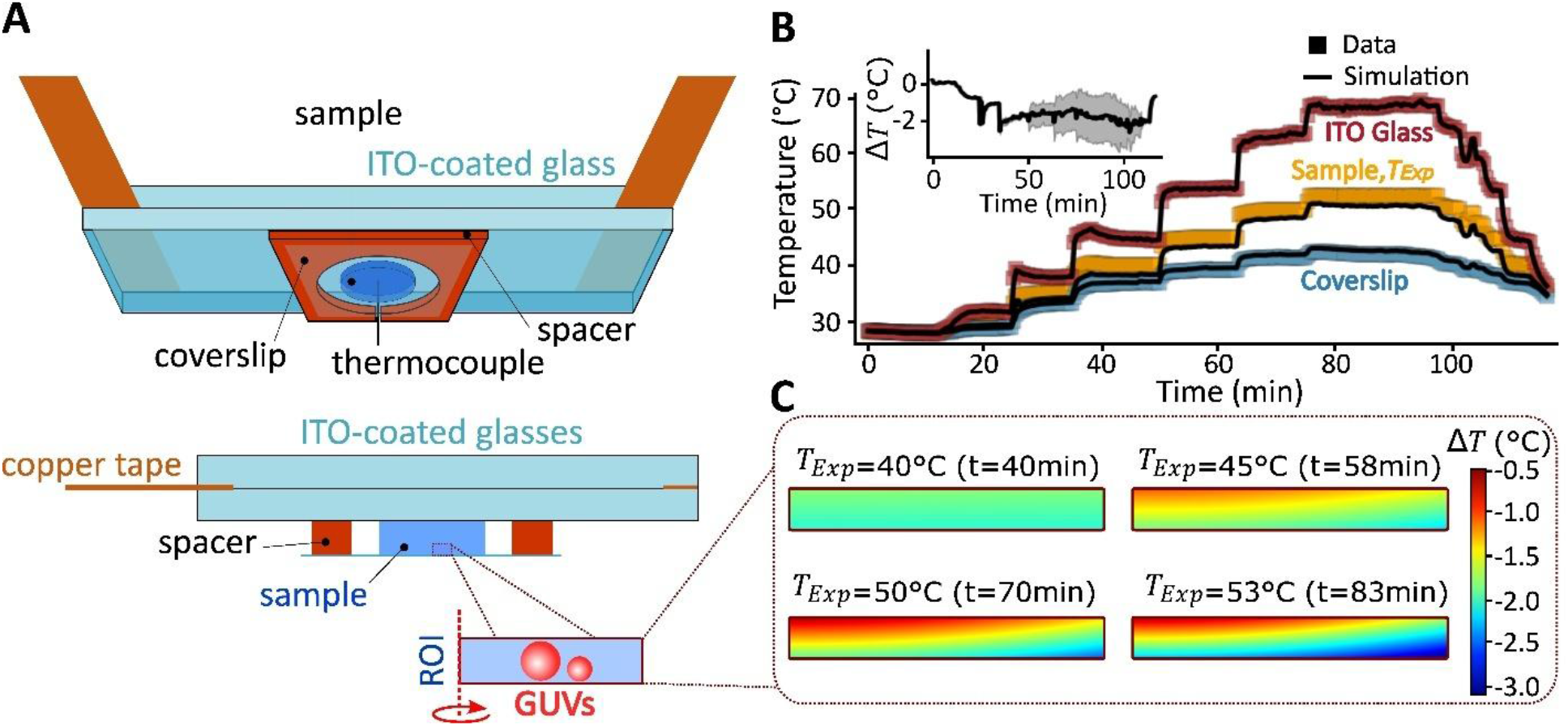
Temperature distribution in the custom confocal heating chamber and its numerical calibration. (A) Schematic of the temperature-controlled device (see Fig. S9 for drawing with dimensions). Top: 3D perspective of the sample and experimental setup. Bottom: 2D cross-section schematically illustrating the heating geometry. The region of interest (ROI) where the microscopy measurements and the simulations were performed is a cylindrical volume of 0.5 mm radius and 0.05 mm height at the base of the chamber (the width of the displayed ROI is not to scale); red spheres schematically represent vesicles of 50 and 20 µm diameter and the red dashed line represents the axis of rotational symmetry. The two additional thermocouples above the ITO glasses and below the sample are not displayed (see Fig. S4 for details). (B) Temperatures measured at the top of the device (red), within the sample droplet (orange) *T*_*Exp*_(*t*), and below the coverslip (blue), shown as squares (experimental) and black solid curves (numerical simulations), as a function of time. The inset shows the difference 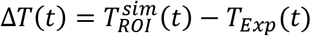 between the average simulated temperature within the ROI, 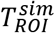, and the thermocouple measurement within the sample, *T*_*Exp*_ ; the shaded region represents the uncertainty based on the maximum and minimum simulated temperatures within the ROI. (C) Simulated temperature distribution within the ROI at different temperature plateaus in panel B corresponding to the following reading of the thermocouple measuring within the sample: *T*_*Exp*_ (*t*=40min) = 40°C, *T*_*Exp*_ (*t*=58min) = 45°C, *T*_*Exp*_ (*t*=1h10min) = 50°C and *T*_*Exp*_(*t*=1h23min) = 53°C. The color scale represents the numerically computed local temperature deviation Δ*T* from the sample thermocouple reading at each plateau. The simulated temperature ranges, 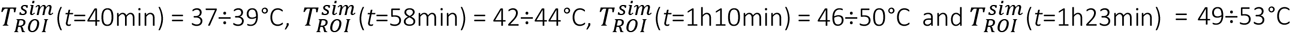, illustrate how the thermal gradient across the ROI, and hence the overall individual vesicle surfaces, evolves with increasing setpoint temperature. Note how the temperature experienced by a GUV can differ substantially from the value reported by the sample thermocouple.

Because heating is applied from above, a vertical temperature gradient develops within the sample, as evidenced by convection currents in the sample that become more pronounced at elevated temperatures. This gradient leads to a systematic deviation between the thermocouple reading and the actual temperature experienced by GUVs resting at the bottom of the chamber (Fig. 4B). This limitation is not unique to our device: in most commercial heating stages, temperature sensors are considerably larger than a vesicle (thus averaging over a larger sample space) and located several hundred microns from the focal plane, while heating is rarely applied homogeneously across the sample.

To estimate the temperature at the GUVs more accurately and assess the magnitude of the temperature gradients, we conducted numerical simulations using thermocouple measurements at two positions: at the ITO glass surface directly above the sample and in the space between the objective and coverslip below the chamber (Fig. 4C, Fig. S4). Throughout this work, we report temperatures as measured by the thermocouple within the sample (hereafter, sample temperature), and accompany each value with the corresponding simulated temperature at the GUV location (hereafter, simulated GUV temperature; note that this represents a range rather than a single value, reflecting the spatial extent of the thermal gradient within the region of interest or ROI), allowing direct comparison across experiments. Our simulations reveal two aspects worth noting at elevated sample temperatures. First, the temperature gradients within the sample are steep enough that a GUV of diameter ∼50 µm experiences a temperature difference of roughly 3°C between its bottom and top (see simulated temperature distribution at a sample temperature of 50°C in Fig. 4C). Second, the temperature gradients across the whole chamber are substantial, resulting in a difference between the sample thermocouple reading and the simulated GUV temperature: at a sample temperature of ∼50°C, the GUV temperature is up to ∼4°C lower, while at ∼40°C this difference reduces to ∼3°C (see inset in Fig. 4B and Fig. 4C). These gradients are specific to our chamber geometry (800 µm height, Fig. S9) and are expected to increase further for thicker sample chambers, making instrument-specific calibration particularly important.

Thermal gradients of several Kelvin can be consequential for experiments in which phase transitions spanning only a few degrees are being investigated by microscopy. In the present study, however, these corrections are modest relative to the expected breadth of the gel-fluid transition in azoPC:DPPC GUVs (section 3.3), and do not alter our qualitative conclusions. Importantly, the combination of our custom heating stage with numerical simulations allows us to reliably estimate the actual temperature at the single-GUV level, as we exploit in the following section.

### 3.5. Temperature-dependent domain reorganization

The observations of the vesicles and the detection of domains over time were done using confocal microscopy with excitation at 633 nm, which ensures that azoPC remains predominantly in its *trans* state during imaging. The GUVs were incrementally heated in steps of 5°C and allowed to equilibrate for several minutes before each observation as done in Fig. 4B. Even when the sample temperature exceeded the melting transition temperature determined by DSC (31.9°C), for example at a sample temperature of 35°C, the dark gel domains persisted for at least 10 minutes, and remained visible even at a sample temperature of 45 °C (Fig. 5A). This persistence of gel domains well above the DSC peak transition temperature is consistent with the expectation of a very broad gel-fluid transition in GUVs discussed in section 3.3, where we argued that coexistence could extend more than 10 K above the calorimetric peak.

**Figure 5.**
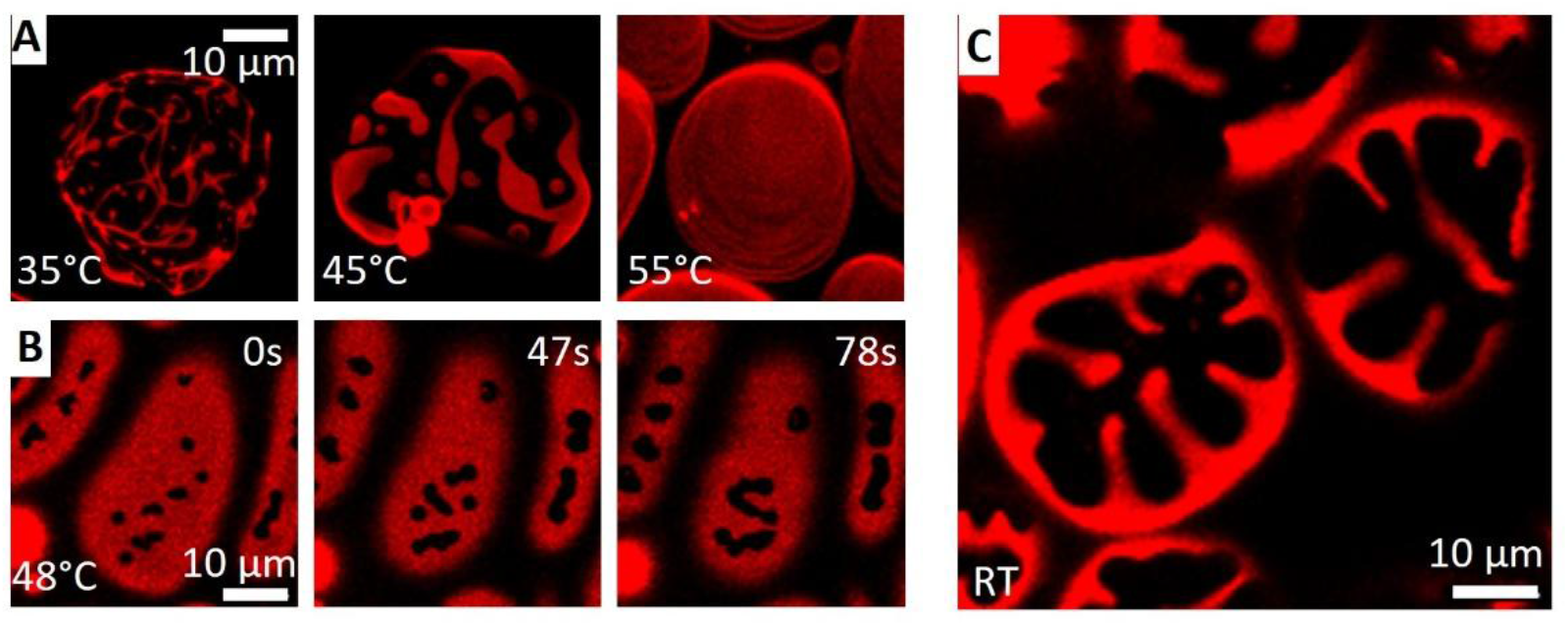
Temperature-dependent melting and domain reorganization in azoPC:DPPC (1:1) GUVs. (A) Domain morphology at successive stages of heating of the same vesicle. Sample temperatures from left to right: 35, 45, 55 °C, corresponding to simulated GUV temperature ranges of 32÷34°C, 42÷44°C, and 51÷55°C respectively (see Fig. 4C for the simulated ROI temperature distribution). Images were acquired after ∼5 min equilibration at each temperature setpoint. (B) Small, circular gel domains nucleate and their boundaries subsequently merge at a fixed sample temperature of 48°C (simulated: 42÷48°C); see Movie S4. Time 0 corresponds to the beginning of the recording (roughly after stabilizing the target temperature). (C) Large flower-like gel domains observed at the bottom of deflated GUVs at room temperature following a controlled heating-cooling cycle in which the sample was first heated until all domains had fully melted, then cooled until the first small domains were observed by microscopy, held at that temperature for approximately 5 min to allow domain coarsening, and finally allowed to cool passively to room temperature. Domain coarsening during the slow cooling step produces the large domains visible here, in contrast to the dendritic patterns that reform under faster continuous cooling (Fig. 1B).

In inverted microscope configurations, where heating is applied above the sample while heat is lost through the objective and coverslip below, thermocouple measurements systematically overestimate the temperature experienced by GUVs resting at the bottom of the chamber. This represents a potential source of discrepancy between DSC-determined transition temperatures and those inferred from microscopy that, to our knowledge, has not been explicitly quantified in the GUV literature. Here, the GUVs only fully melted at sample temperatures well above 50 °C (Fig. 5A, last image), which corresponds to 46÷50 °C as assessed by numerical simulation (Fig. 4B).

Starting from fully melted membranes at 55°C sample temperature (51÷55°C simulated) and during cooling, small, round gel domains reappeared at a sample temperature of around 48 °C (44÷48°C simulated), see Fig. 5B. The appearance of discrete, circular domains is characteristic of nucleation and growth below the binodal boundary, where the circular shape minimizes the line tension energy of the gel-fluid interface. Over time, the domain boundaries appeared to merge while the overall irregular shape of the merged domains was preserved, consistent with arrested coarsening at lowered membrane fluidity at decreasing temperature (Fig. 5B; Movie S4)^49^. Note that these observations were recorded at the bottom of the vesicle, within a region of interest considerably smaller than the full ROI used in the simulations, indicating that the actual temperature experienced by the observed GUVs was close to the lower bound of the simulated temperature range, namely 44°C.

Upon returning to room temperature, the domains further coarsened, with gel domains localizing preferentially at the top and bottom of deflated vesicles, forming large flower-like patterns (Fig. 5C). This spatial organization reflects the tendency of rigid gel domains to avoid regions of high membrane curvature: slightly deflated vesicles adopt an oblate geometry in which the equatorial rim is more curved than the top and bottom caps, and the energetic cost of bending stiff gel domains drives their redistribution toward the flatter regions. This behavior is consistent with theoretical models treating gel domains as rigid crystalline inclusions embedded in a fluid membrane ^50^.

When cooling was performed continuously (rather than stepwise as above allowing for coarsening time), implying an overall faster cooling rate and no equilibration pauses, finger-like dendritic patterns reformed (Fig. 1B) instead of the coarsened domains seen in Fig. 5C. The dendritic morphology is consistent with diffusion-limited domain growth kinetics under rapid cooling, where the system is unable to equilibrate and domains grow anisotropically along directions of fastest lipid transport. In equimolar mixtures, this pattern has been associated with spinodal decomposition^49^, though the distinction from fast nucleation and growth in this composition range warrants careful interpretation.

The microscopy observations confirm that the gel-fluid transition in azoPC:DPPC GUVs is considerably broader than the DSC profile suggests: domains remained detectable approximately 18 °C above the DSC peak transition temperature and approximately 13 °C above the end of the DSC peak baseline (Figs. 3 and 5A), providing direct visual evidence for the extended coexistence region speculated in section 3.3. Additional contributions from lipid oxidation or photobleaching during continuous imaging^51^ cannot be excluded, though the independence of the apparent transition temperature on equilibration time argues against latent heat effects as a primary factor.

Additionally, a small fraction of domains failed to melt fully even at sample temperatures of 50°C (46÷50°C simulated), persisting for at least 5 minutes (Fig. S10). This behavior is most plausibly attributed to two factors. First, large, compact gel domains have a lower perimeter-to-area ratio than smaller ones, meaning that lipid exchange between the gel domain and the surrounding fluid phase is kinetically slower and the domain interior is effectively shielded from the phase boundary and dissolves on a longer timescale even when thermodynamically unstable above T_m_. Indeed, the size dependence of melting kinetics has a straightforward physical analogy: just as an iceberg melts more slowly than a snowball of the same composition due to its smaller surface-to-volume ratio, several-microns-large gel domains in GUVs dissolve more slowly than sub-microscopic (nanometric) domains in MLVs and LUVs. This could be why, at similar heating/cooling rates, bulk calorimetric measurements on those smaller assemblies report a sharper transition than direct microscopy of GUVs. This kinetic trapping of large domains upon heating is the mirror image of the arrested coarsening observed upon cooling discussed above. Second, trace impurities in the lipid preparations are known to stabilize gel domains anomalously, as demonstrated for DPPC GUVs where glucose-derived contaminants produced dye-depleted domains persisting well above the main transition temperature^47^. The combination of these two effects, namely slow dissolution kinetics of large domains and impurity-mediated stabilization, provides a sufficient explanation for the residual domains observed here, without invoking an intrinsic elevation of the transition temperature of the lipid mixture itself.

We also explored the use of LAURDAN as an environmentally sensitive fluorescent reporter of the gel-fluid transition, following its established application in GUV imaging^48^. In that study, LAURDAN two-photon microscopy was used to image coexisting gel and fluid domains in binary DPPC-containing GUVs. This binary system with transition temperatures in the range 15–41 °C depending on composition is directly comparable to our system. LAURDAN reports the gel-fluid transition through a red shift in its emission spectrum upon moving from gel to fluid environments, providing a spatially resolved readout of membrane order. However, LAURDAN turned out not to be suitable as a phase indicator in azoPC:DPPC membranes due to the overlap of its excitation-emission spectrum with the photoisomerization spectrum of azoPC, which gives rise to Förster resonance energy transfer (FRET) between the two molecules. As a consequence, when LAURDAN was present at 0.5 mol% in azoPC-containing GUVs at room temperature, the vesicles exhibited no detectable fluorescence and no detectable respond to UV or blue light irradiation in terms of morphology changes (Fig. S11).

We should note that, potential tension-induced shifts in the observations in this section can be ignored as the vesicles were deflated. Micropipette aspiration experiments have demonstrated that increasing membrane tension decreases the miscibility transition temperature in systems of liquid-ordered/liquid-disordered phase coexistence, with a tension increase of ∼0.1 mN/m producing a shift of several tenths of a Kelvin^52^. While that study addressed liquid-liquid coexistence specifically, the same thermodynamic argument is expected to apply to gel-fluid phase boundaries ^53^, and we therefore emphasize the precaution of working with deflated vesicles.

### 3.6. Photoinduced domain nucleation in fully melted azoPC:DPPC membranes

At temperatures where gel domains have fully melted and the membrane is homogeneous, we exposed the vesicles to UV light. In homogeneous fluid azoPC membranes, UV irradiation is known to produce membrane area expansion stored in transient buds^25^ (see also Fig. S6a), and similar area expansion was observed in fully melted azoPC:DPPC vesicles at a sample temperature of 55°C (51÷55°C simulated). In addition, however, we observed an unexpected phenomenon: small (∼3-6 µm) flower-like, six-lobed dye-excluding domains appeared under UV irradiation, a process fully reversed under blue light (Fig. 6; Movie S5). The reversibility under blue light confirms that domain formation is a direct consequence of photoisomerization and not of UV-induced chemical modification or photooxidation of the membrane.

**Figure 6.**
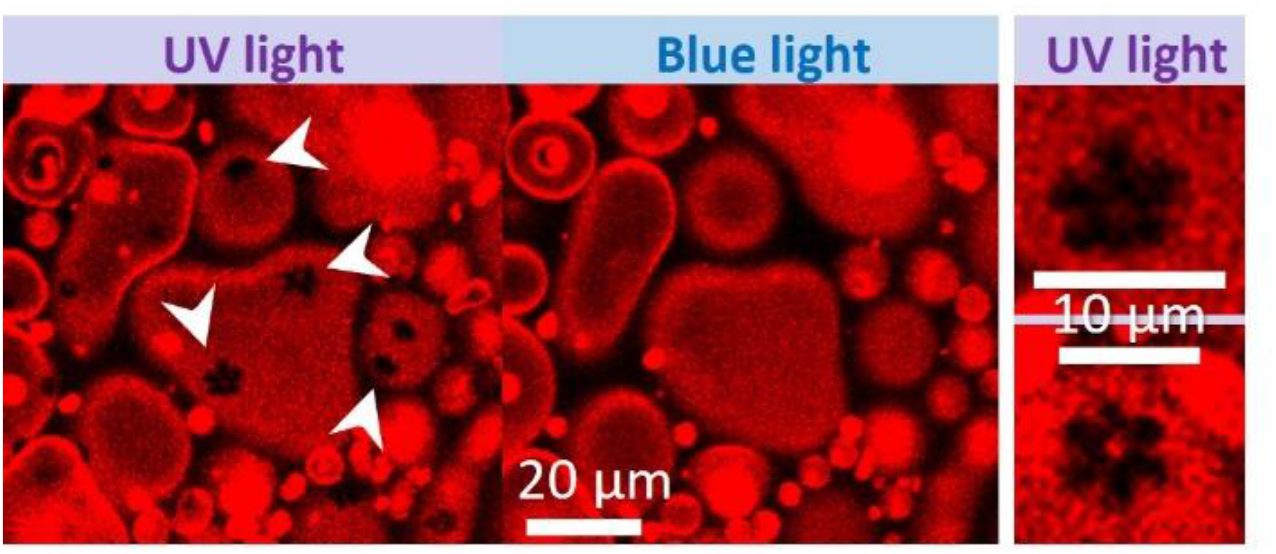
Photoinduced flower-like gel domain formation in fully melted azoPC:DPPC GUVs. Confocal images showing the membrane before UV irradiation (left, homogeneous fluorescence indicating a fully mixed fluid membrane), under UV irradiation (middle, small six-lobed dye-excluding domains indicated by arrowheads), and after blue light exposure (right, full recovery of the homogeneous state); see Movie S5. The flower-like domains, approximately 3-6 µm in size, appear within ∼5s of UV irradiation at ∼74mW/cm^2^ and dissolve fully upon blue light irradiation, confirming their photoisomerization origin. Images were acquired at 55 °C (51÷55 °C simulated).

The unexpected appearance of gel-like domains in a membrane that is fully melted well above the bulk T_m_ of the mixture is puzzling. We considered several candidate mechanisms in turn, noting the deficiencies of each before arriving at the most plausible interpretation.

A first hypothesis is that *cis*-azoPC molecules self-associate via π–π stacking of their azobenzene moieties, locally clustering in a way that excludes the fluorescent dye and mimics a gel-like environment. However, *trans*-to-*cis* isomerization in pure azoPC supported bilayers has been shown to increase lipid diffusivity approximately twofold^45,54^, demonstrating that the *cis* state produces a more fluid, not less fluid, membrane environment. Self-association of azoPC into a gel-like phase is therefore inconsistent with measured molecular mobility data.

A second hypothesis could be related to potential, incontrollable leaflet compositional asymmetry: if azoPC is enriched in one leaflet relative to the other, UV irradiation would generate differential area expansion between the two leaflets, imposing a differential stress that has been shown both experimentally and theoretically to nucleate gel domains in otherwise fluid membranes^55,56^. While this mechanism is physically well-grounded, it is unlikely to be relevant here: electroformation is well established to produce symmetric PC bilayers, and there is no a priori reason to expect preferential partitioning of azoPC to one leaflet during vesicle preparation. Furthermore, upon UV irradiation, potential leaflet asymmetry would have resulted in large spontaneous curvature as leaflet area difference is enhanced by the asymmetric distribution of azoPC triggering tube formation, which we do not observe.

A third hypothesis, and in our view the most physically compelling, is that photoisomerization effectively shifts the phase diagram of the azoPC:DPPC mixture to higher temperatures. In particular, we speculate that the *trans* and *cis* isomers of azoPC have different transition temperatures when mixed with DPPC. While our DSC measurements established a transition temperature of 31.9°C for the *trans*-azoPC:DPPC system (Fig. 3), the geometric incompatibility between the kinked *cis* tail and the straight acyl chains of DPPC (Fig. 1A) may raise this transition temperature substantially upon UV irradiation. The severely kinked tail of *cis*-azoPC makes it a far poorer co-solvent for DPPC in the fluid phase, reducing the free energy of mixing and effectively shifting T_m_ of the mixture upward. If this upward shift is sufficient to bring the effective T_m_ of the *cis*-azoPC:DPPC system above the actual temperature at the GUVs, DPPC-rich gel domains will nucleate spontaneously in the newly supersaturated membrane.

This hypothesis accounts naturally for all key observations. It explains why the domains are dye-excluding DPPC-rich gel domains: DPPC is expelled from the *cis*-azoPC-rich fluid phase as its solubility in it decreases. It explains the contrast with cholesterol-containing liquid-ordered/liquid-disordered systems, where UV light suppresses rather than induces phase separation^31^: in those systems the relevant comparison is between *cis*-azoPC and cholesterol as ordering agents, not between *cis* and *trans* as co-solvents for a gel-forming lipid, and the absence of cholesterol in our system is therefore the critical compositional difference. Finally, it explains the complete reversibility under blue light: isomerization to *trans*-azoPC recovers the more favorable mixing geometry, lowering the effective T_m_ and dissolving the domains. A direct experimental test of this hypothesis would be a DSC measurement on *cis*-azoPC:DPPC multilamellar vesicles, which would be predicted to exhibit a higher transition temperature than the *trans* system characterized in Fig. 3. However, such an experiment is technically challenging (if not unfeasible) due to the spontaneous thermal relaxation of *cis*-azoPC.

The six-lobed morphology of the photoinduced domains, reminiscent of the flower-shaped 2D crystalline domains reported by Wan et al. in DPPC:DOPC GUVs^50^, is consistent with the tendency of such rigid domains embedded in a fluid membrane to adopt developable shapes to minimize strain^50^ (developable shapes are those that can bend along one direction while remaining flat along another, like the petals of a flower wrapping around a sphere, thereby avoiding the large elastic strain that would arise from bending simultaneously in two directions). In Wan et al., the petal instability is driven by elevated membrane tension building up during cooling on large vesicles. However, in our system tension is reduced by UV irradiation, so the petal morphology likely reflects the intrinsic preference of a DPPC-rich crystalline domain for developable shapes, rather than a tension-driven instability.

Regardless of the precise mechanism, the reversible light-controlled nucleation and dissolution of gel domains in a fluid membrane represents a qualitatively distinct phenomenon from the crumpling and mechanical frustration described in earlier sections (Fig. 1), and one that highlights the richness of photoswitchable binary membranes beyond simple area-expansion effects.

## 4. Conclusions

We demonstrated that photoswitchable lipids provide a versatile handle for dynamically controlling the mechanical response of gel-fluid phase-separated model membranes, with the nature of the response determined by the spatial organization of the coexisting phases.

In azoPC:DPPC (1:1) GUVs at room temperature, UV-induced *trans*-to-*cis* isomerization generates excess membrane area that is stored as mechanical deformation. The character of this deformation depends critically on the size and connectivity of the gel domains (Fig. 1), which is in turn set by the thermal history of the sample. When gel domains are small and dispersed (as in vesicles cooled rapidly through the melting transition) the area expansion generates global crumpling distributed across the entire vesicle, reflecting mechanical frustration between expanding fluid regions and a non-percolating gel scaffold. When gel domains are coarsened by slow stepwise cooling (producing a mechanically percolating network) deformation is spatially confined: flat gel facets remain immobile while the fluid azoPC-rich islands between them protrude as localized, reversible buds. In both regimes the deformation is fully reversed by blue light irradiation, confirming that the response is driven by photoisomerization rather than chemical modification. Membrane integrity is preserved throughout, as verified by the absence of leakage of an encapsulated fluorescent dye (Fig. 1E).

Calorimetric measurements on azoPC:DPPC MLV and GUV suspensions establish a gel-fluid transition temperature of 31.9°C for the *trans* system (approximately 10°C below the T_m_ of pure DPPC, Figs. 3, S8) and temperature-controlled confocal microscopy directly visualizes domain melting and re-nucleation (Fig. 5). Direct microscopy observations confirm that the transition in GUVs is far broader than the calorimetric profile suggests, with gel domains persisting well above the DSC peak, a finding we attribute to the reduced interlamellar cooperativity and larger domain size in GUVs compared to MLVs A custom-built heating stage with numerical temperature calibration revealed that thermocouple-based measurements in inverted microscope configurations systematically overestimate the temperature at the GUV location due to vertical thermal gradients (Fig. 4), a source of systematic error in temperature-resolved GUV experiments that has not previously been explicitly quantified.

Perhaps the most unexpected finding is the light-induced nucleation of gel domains in fully melted, homogeneous membranes held well above the bulk transition temperature of the trans mixture (Fig. 6). We propose that this phenomenon arises because *trans*-to-*cis* photoisomerization raises the effective gel-fluid transition temperature of the binary mixture: the kinked *cis* tail of azoPC is geometrically incompatible with the straight acyl chains of DPPC, reducing their mutual miscibility in the fluid phase and driving DPPC-rich gel domains to nucleate. This represents a photoswitchable, fully reversible shift in membrane phase behavior. This finding is a fundamentally different phenomenon from the area-expansion-driven deformation described in prior work on homogeneous azoPC membranes and on Lo/Ld phase-separated systems containing cholesterol^31^. The small flower-like morphology of the photoinduced domains, with six-fold symmetry consistent with the crystalline packing of DPPC, reflects the tendency of rigid gel inclusions to adopt shapes that minimize elastic strain in the surrounding fluid membrane.

These findings establish several principles with broad relevance. First, gel domain architecture controlled through thermal history functions as a structural parameter that programs the spatial distribution of light-driven membrane remodeling, from globally distributed to locally confined. Second, photoswitchable lipids can modulate not only membrane mechanics but also phase equilibria, enabling reversible, light-controlled switching between mixed and phase-separated states without changing composition or temperature. Third, the temperature calibration methodology developed here is of general value for any quantitative GUV experiment involving a heating stage on an inverted microscope.

Looking ahead, the ability to reversibly and non-invasively switch between global and local membrane deformation modes using light opens possibilities for the design of stimuli-responsive synthetic cells (see e.g. Ref. ^57^), membrane-based actuators in soft robotics^58^, and compartmentalized reaction systems where domain boundaries serve as programmable sites for local morphological change.

## Author contributions

R.D. supervised the project. All authors designed the experiments, T.-W.S. performed the experiments and analyzed the data. M.A. assisted with establishing the irradiation and observation protocols. E.S. performed the numerical simulations. T.-W.S., E.S. and R.D. wrote the manuscript. All authors edited the manuscript.

## Notes

The authors declare no conflict of interest

## Funding sources

This work was supported by Germany’s Excellence Strategy, EXC 2008/1 (UniSysCat), Grant 390540038.

## Acknowledgements

TWS acknowledges the support of Klaus Bienert for developing the heating device control software and setup.

## Data availability

The data supporting this article have been included as part of the Supplementary Information.

## Notes

### Competing Interest Statement

The authors have declared no competing interest.

