## Supporting Information for "Light-controlled membrane remodeling in gel-fluid phase-separated giant vesicles using photoswitchable lipids"

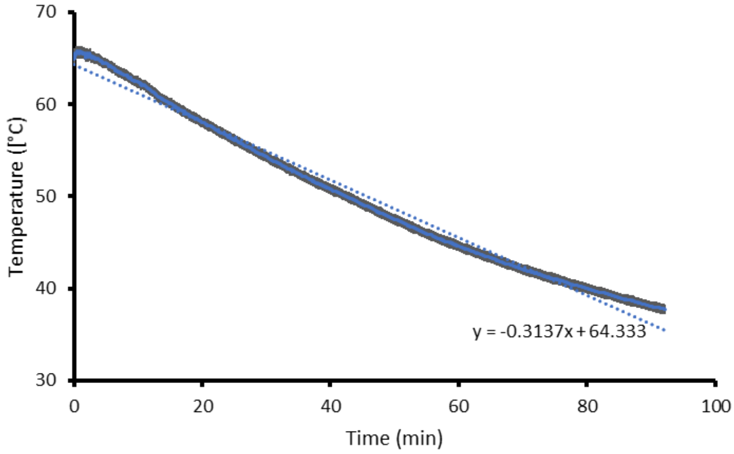


**Figure S1.** Cooling temperature recorded in the oven (where GUVs were electroformed) after switching it off upon completing GUV electroformation. The cooling rate is around 0.3 K/min estimated form the slope of the linear fit (dotted line).

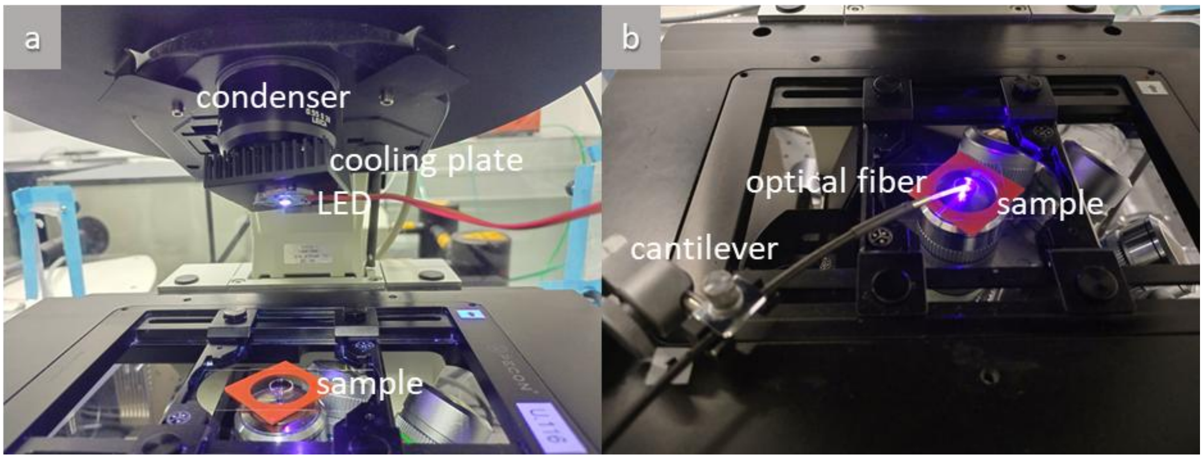


**Figure S2.** Photographs depicting the two approaches used to apply irradiation to the sample: (a) The UV-LED unit is positioned above the sample observation chamber and is attached to the condenser of a microscope (in this picture, the condenser tower is tilted back for better visualization). (b) A fiber-coupled LED system consisting of an optical fiber held by a cantilever over the sample. In both images, the observation chamber is assembled from two coverslips sealed by a Press-to-Seal silicone spacer (red) encircling the sample drop.

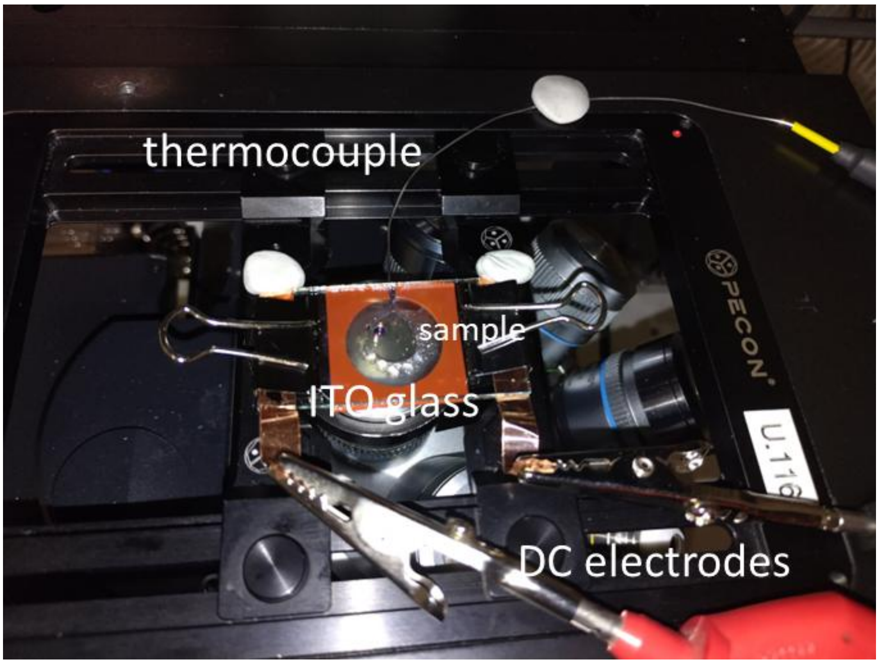


**Figure S3.** Top view photograph of chamber and heating elements for establishing temperature control on GUV samples under the microscope. The detailed sketch of the setup is provided in Fig. S4a in the main text.

### Numerical simulations

Numerical simulations were performed using the finite element method implemented in COMSOL Multiphysics (COMSOL Multiphysics® v. 5.2. www.comsol.com. COMSOL AB, Stockholm, Sweden) on a cluster computer (262 GB RAM, Intel® Xeon® 2.2 GHz (48 CPUs) processor). The model was implemented in a 2D-axisymmetric formulation, as illustrated in Fig. S4a.

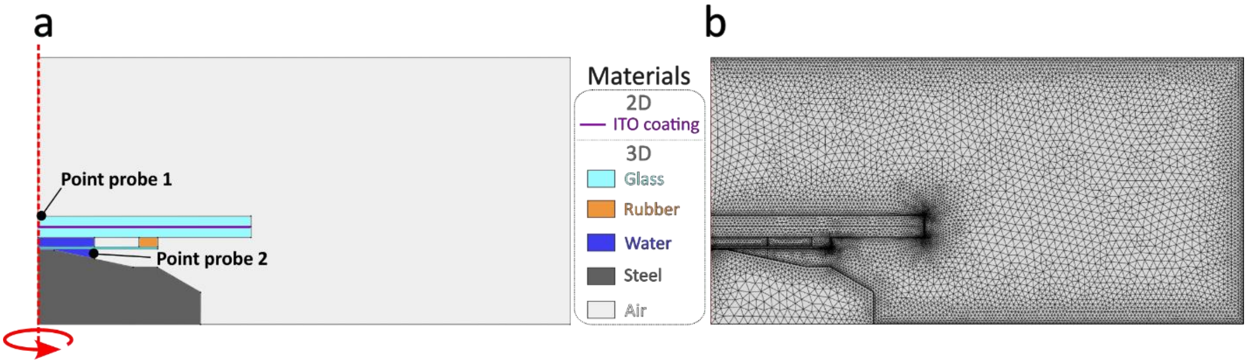


**Figure S4**. Numerical model geometry and mesh. (a) Geometry of the numerical setup. The dimensions of the different parts of the system correspond to those shown in Fig. 4A in the main text and are given in Fig. S9. The red dotted line represents the axis of rotational symmetry. Distinct colored regions represent simulation domains with material properties listed in Table S1. Point probe 1 and point probe 2 refer to the positions at which the two thermocouples were placed (above the top ITO slide and in the space between the objective and coverslip below the chamber) providing values for the respective boundary conditions. (b) Snapshot of the fully meshed computational domain, including refined boundary-layer elements at various interfaces.

The computational domain was discretized using triangular finite elements, with two layers of rectangular boundary-layer elements applied at the fluid interfaces to improve resolution of thermal gradients. A snapshot of the fully meshed geometry is shown in Fig. S4b. The total number of elements in each domain and their corresponding average mesh quality are summarized in Table S1.

In the following, we summarize the sets of equations solved numerically. Values of all material parameters are given in Table S1.
In the fluid domains (water and air in Fig. S4), the following heat transfer equation was solved:

$$\rho C_p \frac{\partial T}{\partial t} + \rho C_p \boldsymbol{u} \cdot \boldsymbol{\nabla} T + \boldsymbol{\nabla} \cdot \mathbf{q} = 0 \ , \tag{S1}$$

where $T$ is the temperature, $\mathbf{q} = -\kappa \boldsymbol{\nabla} \mathrm{T}$ is the heat flux with κ the thermal conductivity, $\boldsymbol{\nabla}$ is the divergence operator, $C_p$ is the specific heat capacity at constant pressure, $\mathbf{u}$ is the velocity and ρ is the density. This equation was coupled to the Navier-Stokes equation under the Boussinesq approximation, such that the density variations were considered only in the buoyancy term $\rho\mathbf{g}$, while the fluid was treated as incompressible, i.e. $\boldsymbol{\nabla} \cdot \boldsymbol{u} = 0$. The momentum equation reads:

$$\rho_0 \left( \frac{\partial \boldsymbol{u}}{\partial t} + \boldsymbol{u} \cdot \boldsymbol{\nabla} \boldsymbol{u} \right) = -\nabla p + \mu \nabla^2 \boldsymbol{u} + \rho_0 \mathbf{g} - \rho_0 \frac{(T - T_0)}{T_0} \mathbf{g} \ , \tag{S2}$$

where $p$ is the hydrostatic pressure, $\mu$ is the fluid viscosity, $\mathbf{g}$ is the gravitational acceleration vector, and $T_0$ is the reference temperature, set to 25°C.
In the solid domains (glass, rubber and steel, see Fig. S4a), the heat transfer was modeled by the heat conduction equation:

$$\rho C_p \frac{\partial T}{\partial t} + \boldsymbol{\nabla} \cdot \mathbf{q} = 0 \tag{S3}$$

The following Dirichlet boundary conditions were imposed at the ITO interface between the two glass slides (purple boundary in Fig. S4a) and on the bottom interface of the steel objective domain:

$$T = T_{input} \tag{S4}$$

The imposed temperature $T_{input}$ was adjusted so that the computed temperatures at the top glass and coverslip bottom positions of the simulated chamber (see point probes 1 and 2 in Fig. S4a) would match the respective experimentally measured temperatures.
Built-in open boundary Dirichlet conditions were imposed at the top, side and bottom air boundaries:

$$T = T_{ref}, \tag{S5}$$

$$-\boldsymbol{n}.\boldsymbol{q} = 0, \tag{S6}$$

where $\boldsymbol{n}$ represents the interface normal vector and $T_{ref}$ is the room temperature (set to 23°C). Additionally, no-slip boundary conditions

$$\boldsymbol{u} = \boldsymbol{0}, \tag{S7}$$

were applied to the boundaries of all liquid domains (air and water).

**Table S1:** Summary of the computational characteristics and material properties for each domain of the numerical model represented in Fig. S4.

| **Material** | **ITO layer** | **Glass** | **Rubber** | **Water** | **Steel** | **Air** |
|---|---|---|---|---|---|---|
| **Number of elements** | 45 | 810 | 87 | 549 | 587 | 6100 |
| **Average element quality** | 1 | 0.92 | 0.96 | 0.58 | 0.96 | 0.84 |
| **Thermal conductivity** $\boldsymbol{\kappa}$ **(W.m.K$^{-1}$)** | 1 | 1.38 | 0.25 | 0.6 | 44.5 | 0.024 |
| **Specific heat capacity** $\boldsymbol{C_p}$ **(J.kg$^{-1}$.K$^{-1}$)** | 341 | 703 | 1802 | 4184 | 475 | 1005 |
| **Density** $\boldsymbol{\rho}$ **(kg.m$^{-3}$)** | 7000 | 2203 | 1200 | 1000 | 7850 | 1.225 |
| **Viscosity** $\boldsymbol{\mu}$ **(J.s.m$^{-3}$)** | N.A. | N.A. | N.A. | $10^{-3}$ | N.A. | $1.8 \times 10^{-5}$ |

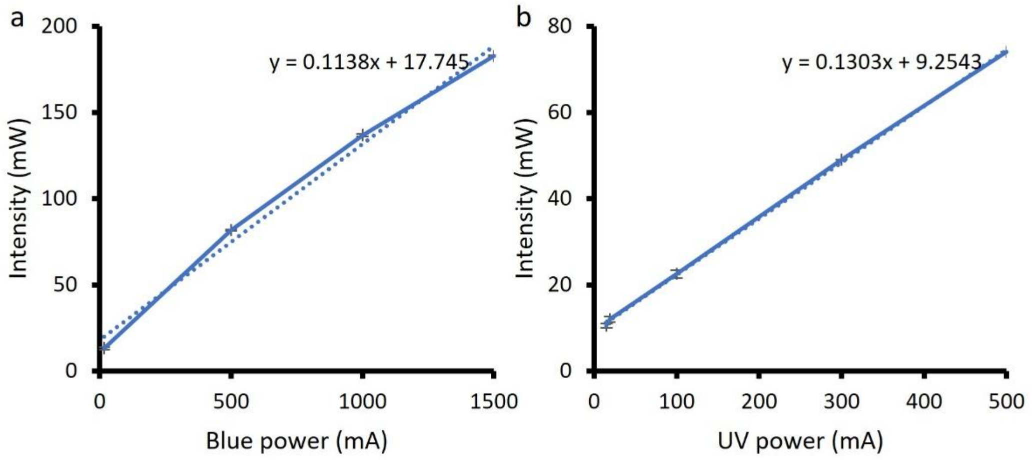


**Figure S5.** Light intensities estimated from (a) blue and (b) UV light fiber-LED device showing the correlation between the setpoint on the device (mA) and the output intensity (mW). The maximum power outputs, measured at the sample plane, were determined using a LaserMate 10 sensor and console (Coherent, USA).

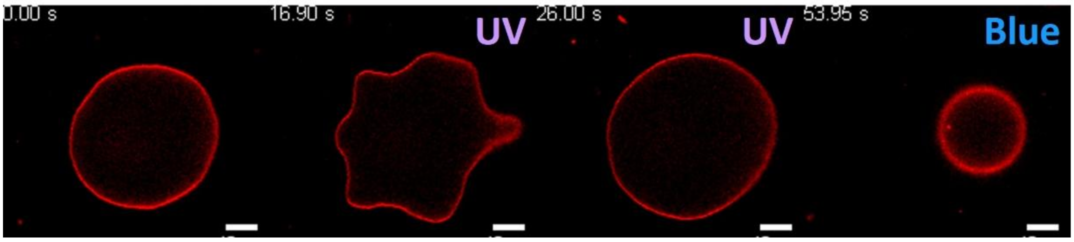


**Figure S6.** Response of fluid azoPC:POPC 1:1 vesicle to UV and blue light irradiation observed under confocal microscopy. The vesicle was irradiated with UV light (~170 mW/cm²) undergoes transient budding (second image) relaxing to a floppier vesicle (third image), a process fully reversed with blue light from confocal laser irradiation (last image). Note that because the vesicle has moved vertically, the confocal cross section shows an apparent smaller diameter under blue light. Scale bars: 10 µm.

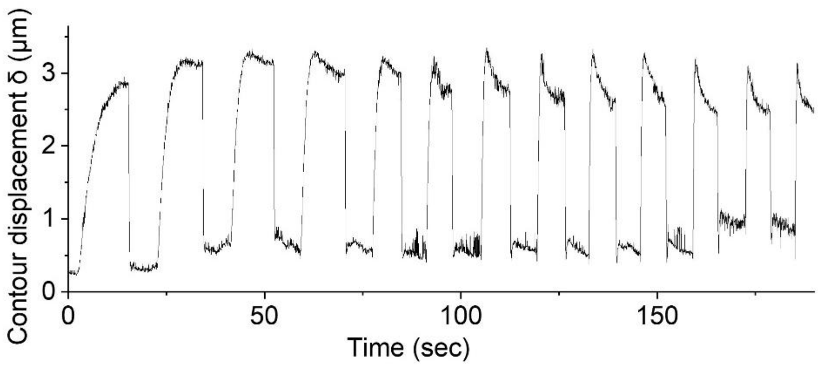


**Figure S7.** Time dependence of the contour deviation (for definition, see Eq. 1 in the main text) of a 1:1 azoPC:DPPC vesicle exposed to UV light (at intensity of 12, 14, 17, 20, 22, 29, 35, 42, 48, 55, 61, 68, and 74 mW/cm$^2$, respectively for the consecutive peaks) and then to blue light (at intensity of 75 mW/cm$^2$) showing fully reversed shape as demonstrated by the example bright field images in Fig. 1B in the main text and Movie S1. The change in the shape of the peaks plausibly reflects slight rotation of the GUV.

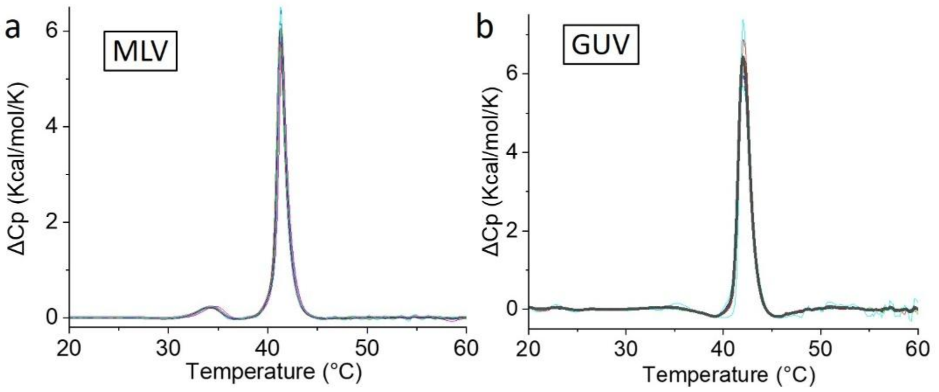


**Figure S8.** DSC heat capacitance curves for DPPC (a) MLVs and (b) GUVs showing main transition peaks at 41.3 °C and 42.0 °C, respectively. The pretransition peak at 34.5°C observed with MLVs is not detected for the GUV sample in which the noise is also stronger. The colored curves represent five consecutive heating scans, with the averaged profile shown as a thick light grey line.

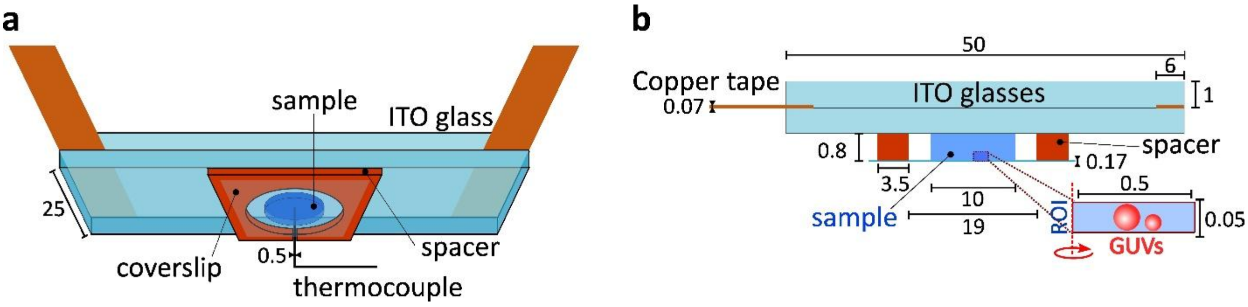


**Figure S9.** Schematic of the temperature-controlled setup in (a) oblique perspective (as shown in Fig. 4 in the main text) and (b) vertical cross-section, the referenced dimensions are in millimeters and are not to scale. The ROI represents the simulated region of interest where the red dashed line represents the axis of rotational symmetry.

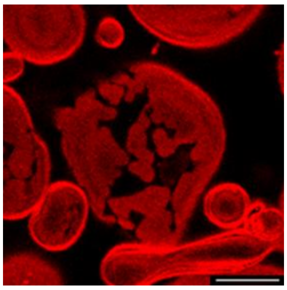

**Figure S10.** Kinetically trapped gel domains during stepwise heating. Confocal fluorescence image of a 1:1 azoPC:DPPC GUV at a sample temperature of 50°C (48±2 simulated). The image was acquired after a stepwise heating protocol (Fig. 5) with approximately 5 minutes of equilibration at each setpoint. Despite the temperature exceeding the main transition, kinetically trapped large gel (dark) domains persist. Scale bar: 10 µm.

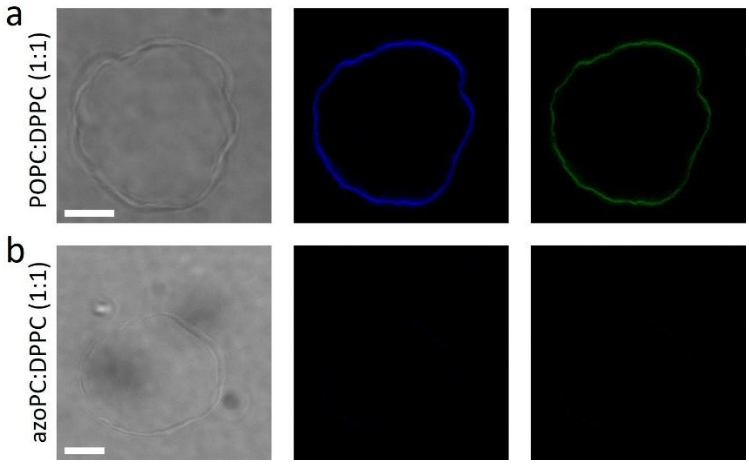


**Figure S11.** Incompatibility of LAURDAN with azoPC-containing membranes. Brightfield and two-photon confocal images of (a) POPC:DPPC (1:1) and (b) azoPC:DPPC (1:1) vesicles containing 0.5 mol% LAURDAN at 23°C. LAURDAN was excited via two-photon excitation at 780 nm. Fluorescence emission was collected in two channels: blue (400–460 nm, middle) and green (470–530 nm, right). In the presence of azoPC (b), LAURDAN fluorescence signal is strongly reduced, likely due to FRET-mediated quenching by the azobenzene moiety. Consistent with this quenching effect, further irradiation with UV or blue light resulted in no detectable morphological changes to the vesicles, indicating a lack of photoswitching-induced membrane area change or LAURDAN-mediated sensitization. Scale bars: 10 µm.

### Movie captions

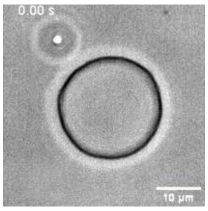


**Movie S1. Reversible global crumpling in a phase-separated 1:1 azoPC:DPPC GUV corresponding to Fig. 1C in the main text.** Brightfield microscopy recording at room temperature. The GUVs were prepared by gradual cooling (~0.3 K/min). Upon UV irradiation (~74 mW/cm²), the vesicle undergoes pronounced global crumpling and twisting deformations distributed across the entire membrane surface, reflecting mechanical frustration between the expanding azoPC-rich fluid domains and the rigid DPPC-rich gel regions that resist uniform area expansion. The crumpled morphology is fully reversed upon blue light irradiation (~75 mW/cm²), restoring the original vesicle shape.

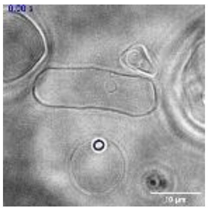

**Movie S2. Reversible localized budding in faceted 1:1 azoPC:DPPC GUVs observed by brightfield microscopy.** Deflated vesicles with coarsened domains at room temperature exhibit flat facets characteristic of rigid DPPC-rich gel regions, which remain immobile throughout irradiation, while the curved fluid segments between them protrude (inward or outward) as localized buds upon UV irradiation (~74 mW/cm²). The buds retract fully upon blue light (~75 mW/cm²), restoring the original faceted morphology. The sequence corresponds to the behavior illustrated in Fig. 1D.

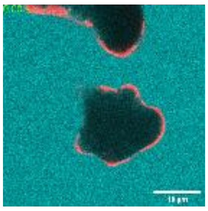

**Movie S3. Reversible localized budding in coarsened azoPC:DPPC GUVs observed by confocal microscopy.** Confocal fluorescence recording of a deflated azoPC:DPPC (1:1) GUV with coarsened gel domains at room temperature. The membrane is labeled with 0.1 mol% Atto-647N-DOPE (red); sulforhodamine B (SRB, cyan) is present in the external solution at ~50 µM. Fluid azoPC-rich domains protrude as localized buds upon UV irradiation (~74 mW/cm²), while the surrounding DPPC-rich gel regions remain flat and unperturbed. Full retraction of the buds is observed upon blue light irradiation (~75 mW/cm²). The absence of SRB signal inside the vesicle throughout confirms that membrane integrity is preserved during repeated deformation cycles. The sequence corresponds to Fig. 1F.

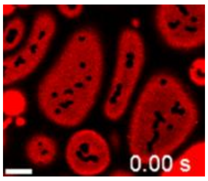


**Movie S4. Nucleation and coalescence of gel domains upon cooling.** Appearance of small circular gel domains at a sample temperature of 48°C (simulated: 44÷48°C) upon cooling, followed by progressive coalescence of domain boundaries. The sequence corresponds to Fig. 5B.

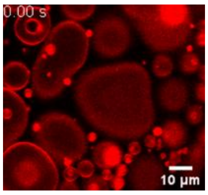


**Movie S5. Reversible photoinduced domain nucleation in a fully melted azoPC:DPPC membrane.** Small six-lobed gel domains nucleate within a homogeneous fluid membrane at 55°C sample temperature (simulated 51÷55°C) upon UV irradiation at 74 mW/cm², and dissolve fully upon subsequent blue light exposure. One also observes slight area increase of the vesicles under UV light. The sequence corresponds to Fig. 6.
